# Late onset of Syt2a expression at synapses relevant to social behavior

**DOI:** 10.1101/2020.01.06.895508

**Authors:** Collette Goode, Mae Voeun, Denver Ncube, Judith Eisen, Philip Washbourne, Alexandra Tallafuss

**Affiliations:** Institute of Neuroscience, University of Oregon, Eugene, OR-97403, USA

## Abstract

As they form, synapses go through various stages of maturation and refinement. These steps are linked to significant changes in synaptic function, potentially resulting in changes in behavior. Here, we examined the distribution of the synaptic vesicle protein Synaptotagmin 2a (Syt2a) during development of the zebrafish nervous system. Syt2a is widely distributed throughout the midbrain and hindbrain early during larval development but very weakly expressed in the forebrain. Later in development, Syt2a expression levels increase, particularly in regions associated with social behavior, and most intriguingly, around the time social behavior becomes apparent. We provide evidence that Syt2a localizes to synapses on socially-relevant neurons in the ventral forebrain, co-localizing with tyrosine hydroxylase, a biosynthetic enzyme in the dopamine pathway. Our results suggest a maturation step for synapses in the forebrain that are spatiotemporally related to social behavior.

## Introduction

During brain development, the assembly of synaptic protein complexes follows regional and temporal signals to form the synaptic connections necessary for a functional network. After intial assembly, synapses go through mulitple stages of structural and functional development, leading to mature synapses that then maintain their final protein compositions over long periods of time (Garner et al., 2006). A multitude of presynaptic proteins are involved in refining highly regulated synaptic properties by controlling the trafficking and release of synaptic vesicles (Rizo and Xu, 2015). Recent studies have added complexity to the maturation process at presynaptic synapses, acquired by switching closely related protein isoforms at already assembled synapses. For example, some synapses undergo a functional Synaptotagmin1 (Syt1) to Syt2 switch during development. This transition has been extensively studied at the calyx of Held synapse in the mammalian auditory brainstem (Kochubey et al., 2016). Development of this large synapse involves several steps during which initial synapses are formed, competing synapses are eliminated and the structure and function of remaining synapses are refined. Fast transmitter release at these synapses initially depends on Syt1, whereas Syt2 has delayed expression onset, replacing Syt1 within a few days, and resulting in faster release kinetics (Kochubey et al., 2016). This example shows the importance of which Synaptotagmin protein is expressed at a synapse, as it determines the synaptic properties.

Synaptotagmins are members of a large family of evolutionarily-conserved, membrane-trafficking proteins, many of which are considered calcium ion sensors that control neurotransmitter release through the SNARE (SNAP Receptor) complex (Südhof, 2002). The large number of Synaptotagmins, with 17 *Synaptotagmin* genes in mammals and 28 in zebrafish, provides the potential to fine tune presynaptic sensitivity and response properties. Often several different Synaptotagmins are expressed in the same tissue and appear to act redundantly after genetic elimination of one protein (Bouhours et al., 2017). However, the specific function of many *Synaptotagmin* genes remains elusive and the need for maintaining this large number of genes is speculative.

Knowing the distribution of Syt2 might help to determine the function it plays during development of specific behaviors. In mouse, Syt2 is highly expressed in the cerebellum, hindbrain and spinal cord. Distinct subsets of neurons express Syt2 in the striatum, *zona incerta*, reticular nucleus of the thalamus and ventromedial nucleus of the hypothalamus at postnatal day 14 (P14) - P16. A developmental change in the Syt2 distribution was reported during the first two postnatal weeks in the mouse retina (Fox and Sanes, 2007), where Syt2 distribution either increased or decreased over time, depending on the specific cell type. Although Syt2 is mostly expressed in inhibitory neurons in the mouse forebrain, Syt2 can be present at both inhibitory and excitatory synapses, even within the same tissue (Chen et al., 2017; Fox and Sanes, 2007; Pang et al., 2006a).

We investigated the expression profile of Syt2 during various stages of zebrafish larval development and, in particular, with a focus on synaptic protein changes that occur at critical time periods, for instance, during the establishment of complex behaviors. We recently demonstrated a requirement for the ventral telencephalon for social behavior (Stednitz et al., 2018), and a number of studies indicate that this behavior initiates around the second week of life (Dreosti et al., 2015; Larsch and Baier, 2018). These observations raise the question of whether synaptic connections of the putative social circuit already exist prior to 14 days postfertilization (dpf), and if they do, what changed to allow the behavior to become functional.

Here we describe a significant increase of the presynaptic protein Syt2a in the zebrafish forebrain during the second week of larval development, compared to relatively steady levels of other pre- and post-synaptic proteins. Syt2a first shows an initially confined forebrain expression restricted to the olfactory bulb, posterior dorsal telencephalon and ventral forebrain region, which then expands to broad and strong expression throughout the forebrain, showing a similar distribution to other synaptic markers. We found that the increase of forebrain Syt2a expression coincides with the temporal onset of social behavior. Further, we show that Syt2a and the catecholamine synthetic enzyme tyrosine hydroxylase (TH) co-localize on projections within the forebrain and on projections originating from dopaminergic cells located in the diencephalon, both of which are important for social behaviors (Alger et al., 2011).

## RESULTS

### Syt2a distribution during early larval development

We examined the protein expression pattern of the presynaptic proteins Syt2a, Synaptic vesicle protein 2 (SV2), Synapsin1/2 (Syn1/2) and the postsynaptic proteins Glutamate receptor 2/3 (GluR2/3) and Gephyrin (Geph) in the brain of zebrafish larvae at 6 dpf (Fig.1A,B). During larval development from preflexion to late flexion stages (6-14 dpf, (Engeszer et al., 2007), immunolabeling for Syt2a was strong in the midbrain and hindbrain, suggesting that Syt2a expression is established early in these brain regions. Contrary to other synaptic markers described here, that showed uniform distribution throughout the brain, Syt2a labeling was weak in the forebrain region (Fig.1A,B, asterisk) at 6 dpf. The forebrain consists of (1) the telencephalon, which processes sensory information, controls motor function and is the most important region for high order function, and (2) the diencephalon, that relays sensory information, influences sensory perception and connects the endocrine and nervous systems. The teleost ventral telencephalon is considered the homolog of the septal area, a part of the limbic system in mammals (O’Connell and Hofmann, 2012). The weak Syt2a signal in the forebrain cannot be explained by weak expression of *syt2a* transcript, as we analyzed *syt2a* mRNA distribution and found that *syt2a* transcript was expressed in the forebrain at 6 dpf in comparable amounts to other brain regions (suppl. Fig. 1). We conclude that *syt2a* transcript is not translated or translated only weakly, resulting in low protein levels in the forebrain at 6 dpf.

**Fig.1.**
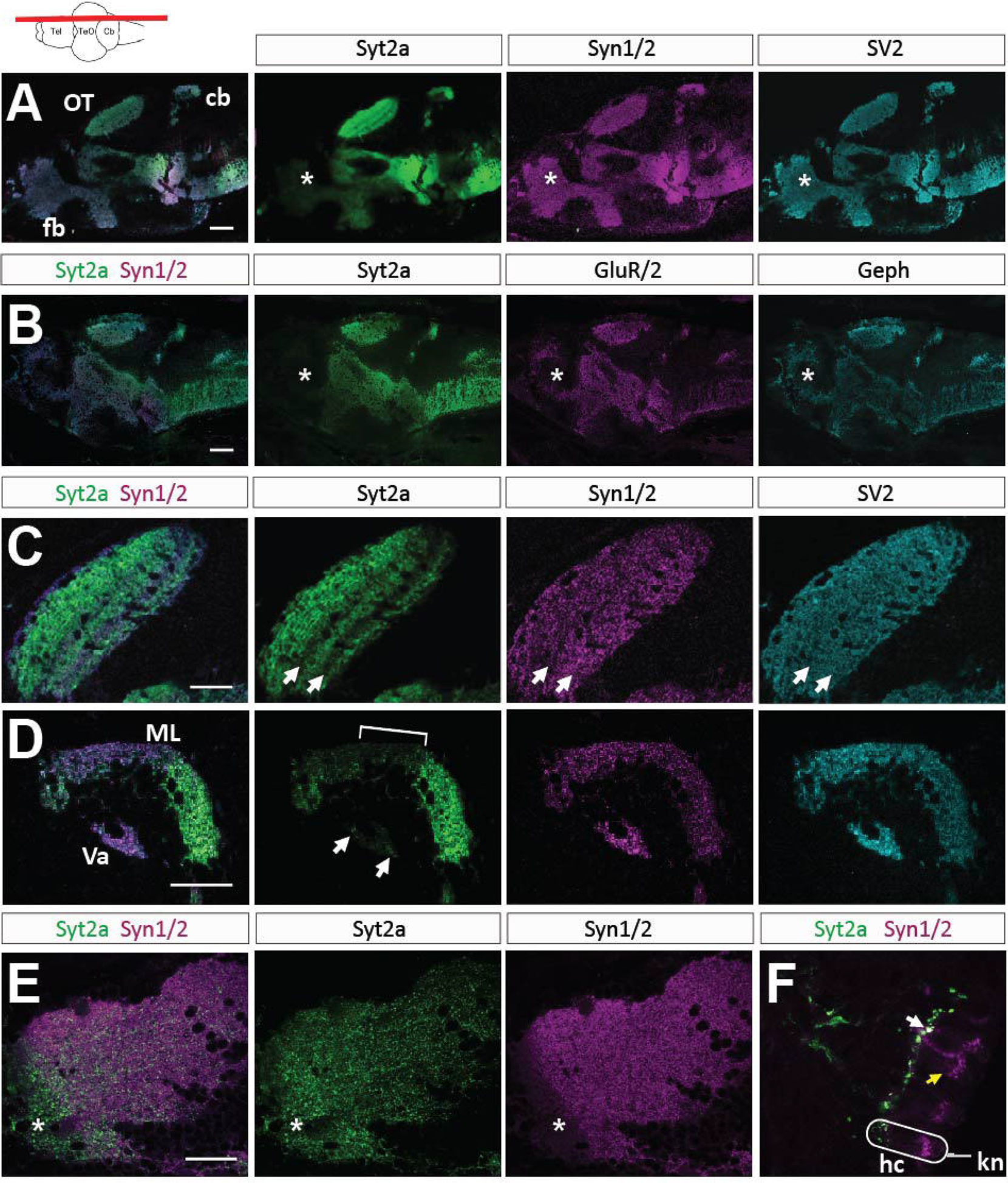
Synaptic distribution of presynaptic proteins in the 6 dpf zebrafish brain. **1A,B**. Overlay and images of individual channels revealing the expression pattern of Syt2, Syn1/2, SV2, Geph and GluR2/3. The forebrain (fb) is marked by asterisks. **1C**. Higher magnification showing individual expression patterns of Syt2, Syn1/2 and SV2. Two arrows mark the low Syt2 labeling in two synaptic layers. **1D**. High magnification of the cerebellum shows Syt2 labeling in the molecular layer (ML), but weaker signal in the medial part (bracket). Arrows point to punctate labeling in the *Valvula cerebelli* (Va). **1E**. The anterior part of the preoptic region (asterisk) shows Syt2 labeling (green) but reduced Syn1/2 labeling (magenta). **1F**. In the lateral line Syt2 is located in projections while Syn1/2 is found in hair cells (hc). White arrow points to co-labeled synapses at the basal face of a hair cell. Yellow arrow points to Syn1/2 labeling in a hair cell. For orientation a single hair cell (hc) and a kinocilia (kn) are outlined in white. All sections except 1F are oriented anterior to the left, dorsal up. The level of the sections are depicted in a cartoon showing a dorsal view of a zebrafish brain (**1A-E**). Scale bars are 100 μm in A, B, 30 μm in C-E and 15 μm in F.

In addition to the distinct weak forebrain labeling, we found other regional differences in Syt2a localization compared to other synaptic proteins. We detected layered Syt2a labeling in the optic tectum (OT) neuropil, with accumulation of Syt2a protein in the fiber-rich layers (Fig.1C, arrows point to layers with weaker Syt2a labeling). Syn1/2 and SV2 did not show this layered pattern, but instead presented an even distribution throughout the OT neuropil. Further, Syt2a was absent from the outermost layer of the OT, labeled by Syn1/2 and SV2. In the cerebellum (Cb), we found regionally distinct Syt2a distribution in the molecular layer (ML, Fig.1D), with sparse labeling in the medial portion of the Cb and strong labeling in the anterior and posterior Cb (Fig.1D, bracket marks sparse labeling in medial portion). Further, Syt2a labeling was weak in the *Valvula cerebelli* (Va, Fig.1D, arrows point to Syt2a puncta).

Although we found that Syt2a was often more restricted in its distribution as compared to Syn1/2, we found one exception in the anterior part of the preoptic area (POA), in which Syt2a showed broad and dense labeling throughout the entire POA, whereas Syn1/2 was absent (Fig.1E, asterisk). Further, in the lateral line neuromasts Syt2a was restricted to projections, whereas Syn1/2 puncta were distributed within the hair cells and on the projections (Fig.1F). Syt2a and Syn1/2 colocalized at the basal end of the sensory receptors, where the projections (green) synapse onto hair cells (magenta, Fig. 1F, white arrow). In summary, Syt2a is more selectively localized in the zebrafish nervous system at 6 dpf, as compared to other synaptic proteins, with the most apparent difference being the very sparse labeling in the forebrain (Fig.1).

### Increase of Syt2a labeling in the forebrain during late larval development

We found that Syt2a labeling in the forebrain increased with development (Fig.2). Whereas Syt2a was barely detectable in the forebrain at 6 dpf, Syt2a labeling increased noticeably at 14 dpf, until it reached an intensity at 18-20 dpf that was comparable to Syt2a expression level in the midbrain and hindbrain (Fig.2A). Despite the low level of labeling at 6 dpf, higher magnification images of the forebrain unveiled sparse, punctate Syt2a labeling in the olfactory bulb (olf), the anterior commissure (AC) and a small region of the dorsal telencephalon (not shown). The AC is a dense neuropil through which many projections of the telencephalon travel and that presents almost no cell bodies. Between 11 and 14 dpf, prominent Syt2a labeling in the ventral telencephalon broadened and expanded to regions adjacent to the initially weak Syt2a labeling. Furthermore, Syt2a signal became more propagated in the olfactory bulb (olf) and within olf projections at 14 dpf (Fig.2B, arrows). At 18 dpf, Syt2a labeled most of the ventral telencephalon and Syt2a expanded in the dorsal telencephalon (Fig.2B, C). We also compared the spatiotemporal distribution of Syt2a and Syn1/2 during forebrain development. As expected, we observed strong and uniform Syn1/2 labeling throughout the forebrain at 6 dpf (Fig.2A-C) with no noticeable change in its distribution or signal strength at 11, 14 or 18 dpf (Fig.2A). These results suggest that although many synapses are already present within the forebrain at 6 dpf, as evidenced by Syn1/2 labeling, most of these synapses do not contain Syt2a until around 14 dpf, a time point at which social behavior is established (Dreosti et al., 2015).

**Fig.2.**
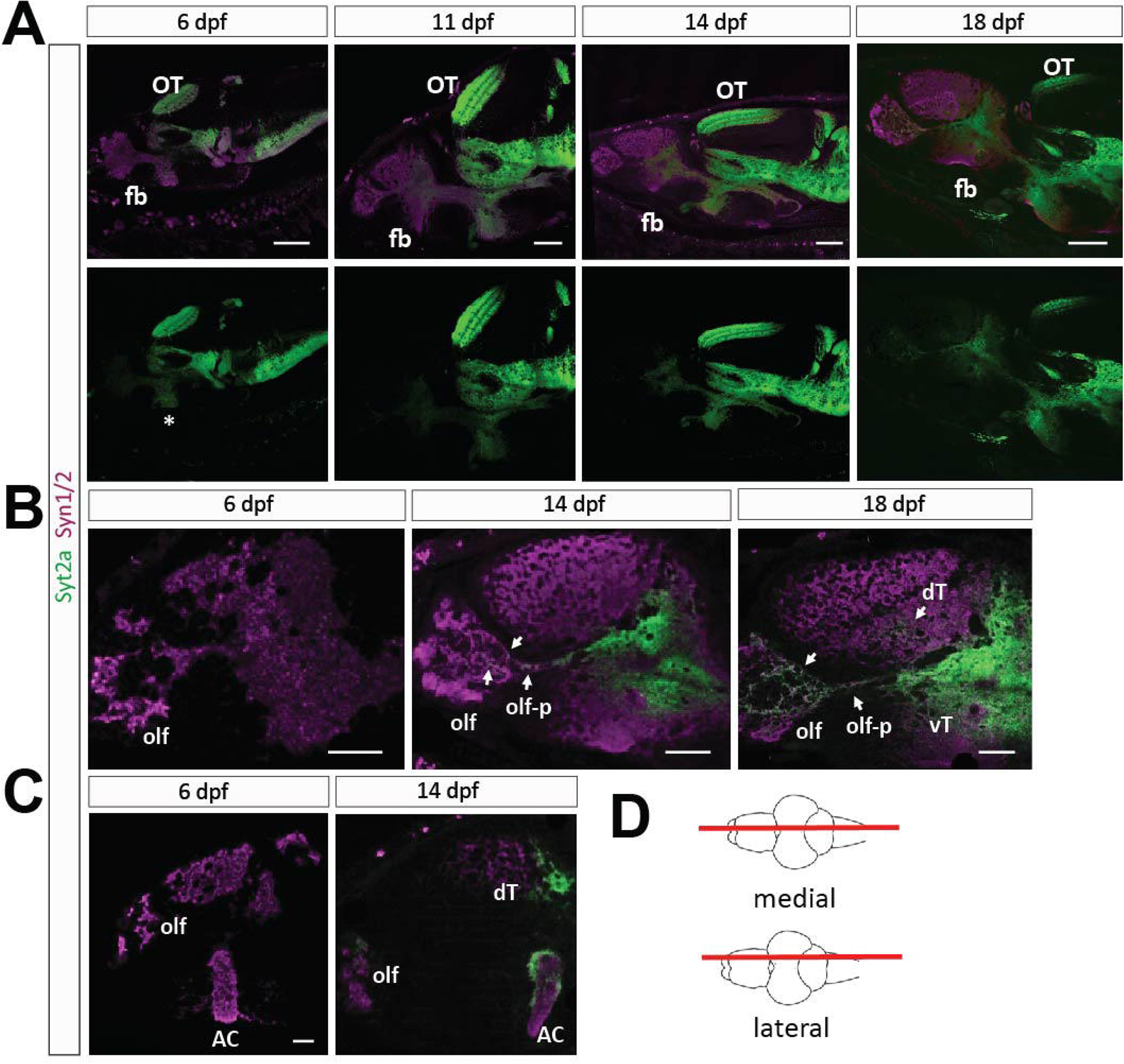
Differences in Syt2a and Syn1/2 labeling during development. **2A**. Sections showing Syt2a and Syn1/2 labeling in the brains of 6, 11, 14, and 18 dpf zebrafish larvae shows that Syn1/2 labeling (magenta) throughout the forebrain (fb) does not change. In contrast, Syt2 labeling (green) becomes more prominent over time selectively in the forebrain. Level of Syt2a labeling in the optic tectum (OT) remains similar among different time points. Medial section (see cartoon in 2D). **2B**. Higher magnification of the forebrain region reveals robust and strong Syt2a labeling in the olfactory bulb (olf), its projections (olf-p) and in the ventral telencephalon (vT) at 14 dpf, which broadens and becomes more visible in the dorsal telencephalon (dT) at 18 dpf. Arrows point to Syt2a puncta within these regions. **2C**. Lateral sections (see 2D) reveal an increase in Syt2a signal in the olf, dT and the dorsal and lateral area of the anterior commissure (AC) between 6 and 14 dpf. **2D**. Cartoon showing a dorsal view of a zebrafish brain marking the level of the lateral and medial sections shown in the images with a red line. Scale bars are 100 μm in A, 50 μm in B and 30 μm in C.

The increased distribution of Syt2a during development is particularly evident in medial telencephalon sections (Fig.2C,D). Syt2a labeling is present in the AC, the olfactory region (olf) and the dorsal telencephalon (dT) at 14 dpf. However, at 6 dpf, Syt2a labeling is almost undetectable in these regions (Fig.2C). We note that Syt2a labeling in the dorsal aspect and on the edges of the AC was denser and appeared stronger compared to other regions within the AC (Fig.2C). We conclude that Syt2a is dynamically expressed in the forebrain during larval development, presumably through regulation of translation, because we did not see dramatic differences in mRNA levels across the brain (Fig.S1).

### Synapse distribution in the anterior commissure shows dynamic developmental changes

We quantified synaptic protein distribution in three selected regions of the forebrain (Fig.3A,C-C”) and found that the relative intensity of synaptic protein labeling changed over development. We analyzed the intensity of Syt2a, Syn1/2 and Geph in the dorso-medial part of the neuropil in the ventral telencephalon (AC, Fig.3B, white square), a region in the posterior part of the dorsal telencephalon (dT) and the anterior part of the hypothalamus (Hy, Fig.3A). We chose three time points, 6, 14 and 20 dpf that reflect different maturation states of zebrafish development. We found the most dramatic increase in Syt2a intensity in the AC and dT between 6 and 14 dpf (Fig. 3C’), where Syt2a intensity peaks at 14 dpf (Fig. 3C-C”, green). In contrast, the intensity of Syt2a labeling in the Hy remained at a similar level throughout this developmental period (Fig. 3C’).

**Fig.3.**
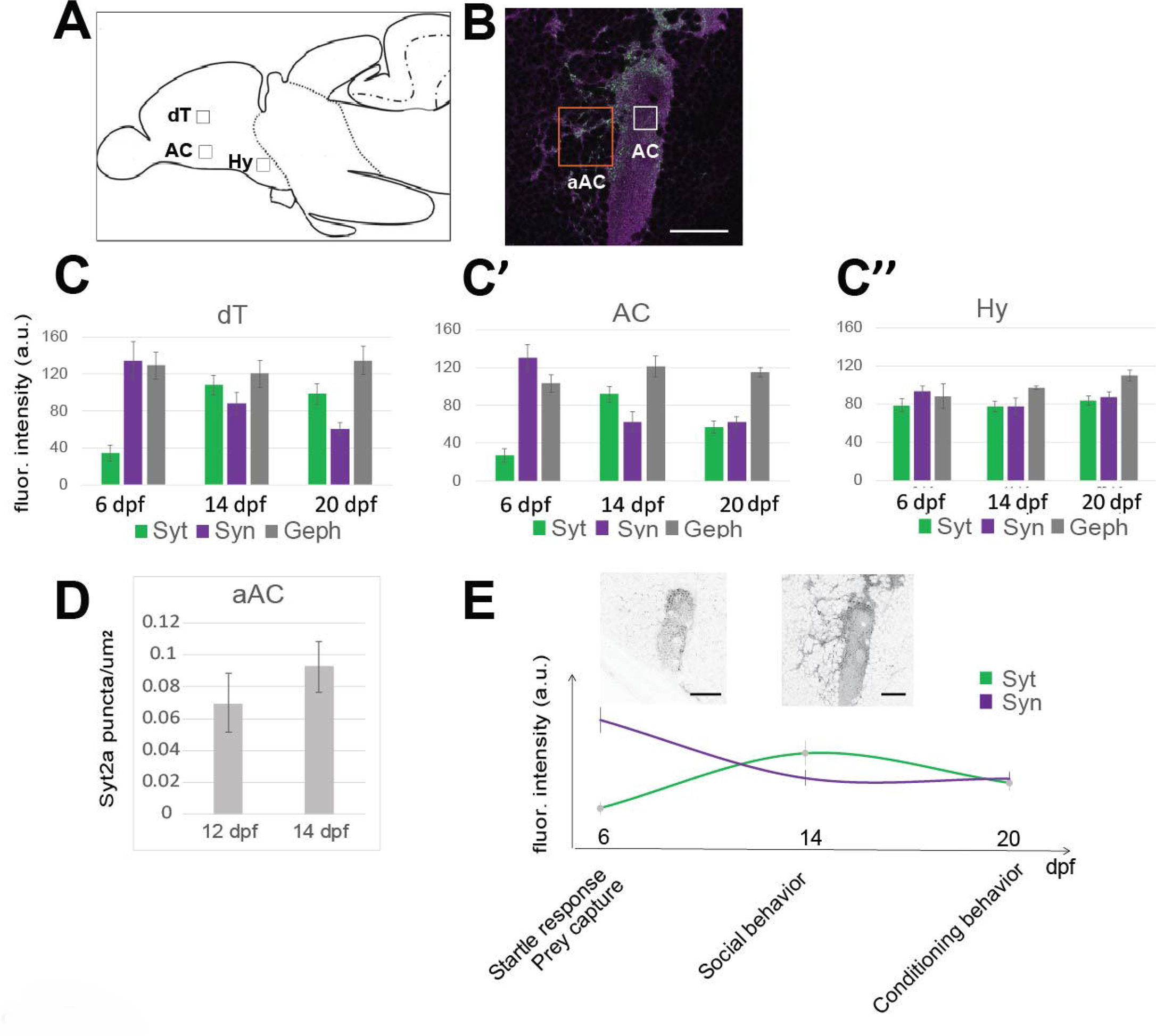
Syt2a expression levels in the telencephalon change over time. **3A**. Cartoon showing a sagittal view of a zebrafish brain. Boxes mark the selected region of interest (ROI) used for synapse counts. **3B**. Image showing the AC labeled for Syt2 (green) and Syn1/2 (magenta) at 14 dpf. The white box shows the actual size of ROI used for counting. The orange box shows an ROI for a synapse count on neuronal cell bodies located adjacent to the AC (aAC). **3C-C”**. Changes in intensity of Syt2, Syn1/2 and Geph labeling at 6, 14, and 20 dpf in the dorsal telencephalon (dT, **3C**), anterior commissure (AC, **3C’**) and hypothalamus (hy, **3C”**). Number of intensity represented in arbitrary units (a.u.). **3D**. The number of Syt2 puncta in the area adjacent to the AC (see 3B, orange box), increases between 12 dpf and 14 dpf. **3E**. Syt2 graph shows Syt2a and Syn1/2 intensity in the AC (see 3C-C”) over time while marking behavioral milestones during zebrafish larval development. The two images show representative Syt2a expression at 6 dpf and 14 dpf, respectively, demonstrating the increase in Syt2 puncta aAC. Scale bars are 30 μm in B and 10 μm in E.

In contrast to Syt2a labeling that is initially low and increases over development, we found that Syn1/2 intensity is already high at the earliest time point tested (Fig. 3C-C”, purple). After peaking at a distinct development stage, the intensity of the presynaptic proteins Syt2a and Syn1/2 finally decreased at a later stage. In contrast, we found that the number of the postsynaptic Geph puncta is stable across the three time points analyzed (Fig. 3C-C”, grey).

We cannot exclude that Geph puncta would increase or decrease at a later time point. We conclude that the dynamic Syt2a expression levels in the telencephalon during development are consistent with process of more synapses acquiring Syt2, whereas the synaptic localization of other synaptic proteins remains relatively stable over the same developmental period.

Consistent with the above observations, we noticed an increase in Syt2a puncta in a region adjacent to the AC (aAC, Fig.3B, orange box), a region populated by neurons which are necessary for social behavior in the adult (Stednitz et al., 2018) and at 14 dpf (AT and PW, unpublished observation). We compared Syt2a puncta at 12 and 14 dpf, a time period in which social behavior becomes more robust, and found a 20% increase in Syt2 labeling in this aAC region (Fig.3D). It is striking that the number of Syt2a puncta becomes prominent and peaks at 14 dpf, a time at which social behavior is significantly strengthened (Fig.3E). These observations suggest that, during this time interval, synaptic properties are modulated, coinciding with the establishment of complex behaviors, including social behavior (Fig.3E) (Dreosti et al., 2015; Larsch and Baier, 2018; Roberts et al., 2013).

### Syt2a-positive synapses are localized on both somata and neurites of forebrain neurons relevant for social behavior

Both inter-fish related social attraction and orienting as well as the detection of biological motion become established around 14 dpf (Dreosti et al., 2015; Larsch and Baier, 2018). We previously identified a genetically-defined population of *lhx8a*-positive neurons in the ventral forebrain of zebrafish larvae that is important for social behavior at 14 dpf (AT, PW, unpublished), as defined by ablation experiments similar to those in adult zebrafish (Stednitz et al., 2018). These cells are cholinergic (Stednitz et al., 2018), similarly to a population of cells in the basal forebrain in mammals (Zhao et al., 2003). Despite the identification of these homologous neuronal populations in both fish and mammals, we know little about the neurons that synapse onto these basal cholinergic forebrain neurons (bCFNs) or what brain regions these neurons innervate.

As a marker for bCFNs, we took advantage of the transgenic line *Et(REX2-SCP1:GAL4FF)y321;Tg(UAS:GFP)*, an enhancer trap located in the *lhx8a* gene, and analyzed Syt2a distribution on these socially-relevant neurons (SRNs) and their projections at 14 dpf. The transgenic line labels two populations of neurons in the forebrain, one in the ventral telencephalon and another in the anterior part of the diencephalon, the preoptic area of the hypothalamus (Hy, Fig.4A). The neuronal population in the telencephalon consists of two clusters, divided by the AC (Fig.A,4B). As described earlier, Syt2a puncta were distributed throughout neurons adjacent to the AC (Fig.4C-C”). Although Syn1/2 showed an even distribution throughout the medial AC (medAC, Fig. 4B,C), we found that Syt2a accumulates at the dorsal aspect of the medAC and on the external face of the neuropil (Fig.4D), whereas the center and the ventral neuropil appears less densely populated with Syt2a labeling. In the region anterior to the AC, we found Syt2a puncta localized at the cell bodies of bCFNs (Fig.4C-C”, arrows). All these puncta were also positive for Syn1/2, suggesting that these are *bona fide* presynaptic terminals (4C”). A few days later, at 20 dpf, we found bCFNs in close proximity (arrows) or even engulfed (asterisk) by Syt2a-positive AC neuropil (Fig.4D,D’), suggesting that Syt2a-positive AC neuropil dimensions expand over time. Another possibility is that more bCFNs form in close apposition to the already established neuropil. We conclude that bCFNs are in close proximity to the synaptic neuropil of the AC, and that Syt2a-positive presynaptic terminals localize to the somata of bCFNs in the ventral telencephalon.

**Fig.4.**
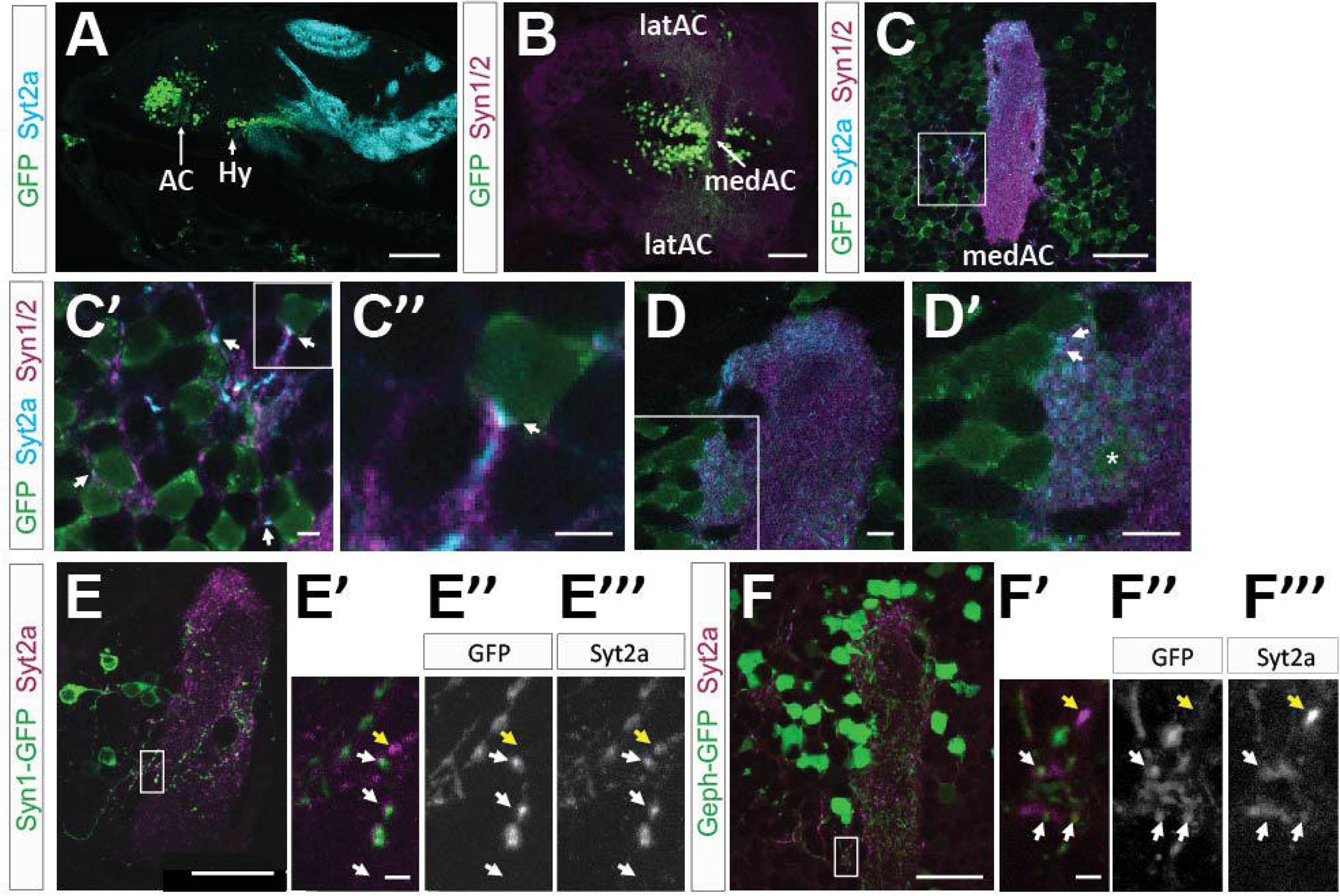
Syt2 labeling in bCFNs and their neurites. **4A**. Sagittal view of Syt2 expression (cyan) in a transgenic fish expressing GFP under control of the y321 enhancer trap, labeling bCFNs. In the forebrain, the largest cluster of GFP-positive neurons are located in the ventral telencephalon, in a region adjacent to the AC, and the anterior hypothalamus (Hy). **4B**. Dorsal view of the forebrain showing a cluster of GFP-positive neurons adjacent to the AC, and divided by the medial AC (medAC). **4C**. Lateral view of the medAC showing Syt2 and Syn1/2 within the GFP-positive bCFN domain. **4C’**. Magnification of the area marked by the box in 4C. Arrows mark presynaptic terminals onto GFP expressing bCFNs. **4C”**. Single neuron, see box in 4C’, with Syt2 and Syn1/2 puncta on the neuronal soma (arrow). **4D**. Higher magnification of the dorsal part of the medAC, showing the close proximity of GFP-positive neurons to the medAC at 20 dpf. 4D’. High magnification with a asterisk marking a GFP-positive neuron enclosed within the medAC neuropil. Arrows mark Syt2 and Syn1/2 positive puncta on GFP positive neurons. **4E-E”’**. Mosaic expression of Syn1-GFP in bCFN neurons and their neurites in the AC at 14 dpf. **4E’-E”’**. High magnification shows co-localization of Syn1-GFP and Syt2a, marked by white arrows, a GFP negative Syt2a punctum is marked by a yellow arrow. Box in 4E shows the magnified area. **4F-F”’**. Labeling of Syt2 with Geph-GFP fusion protein in bCFNs. High magnification of the boxed in area shows co-localization of Syt2a and Geph-GFP, marked by white arrows. A yellow arrow marks a GFP negative punctum. Scale bars are 100 μm in A, 60 μm in B, 30 μm in C, 2 μm in C’,C”, 4 μm in D,D’, 30 μm in E,F and 5 μm in E’,E”,F’,F”.

Because the AC neuropil is very dense and neurites from all bCFNs are hard to follow, we expressed a Synapsin1-GFP fusion protein in a mosaic fashion using the transgenic line *Et(REX2-SCP1:GAL4FF)y321;Tg(UAS:Syn1-GFP)*. This approach allowed us to follow individual neurites from bCFNs and examine colocalization of Syt2a with zebrafish Synapsin1-GFP from a subset of bCFNs in the medAC (Fig. 4E-E”’). We found colocalization of Syt2a with presynaptic Syn1-GFP puncta, suggesting that Syt2a is located in the presynaptic terminals of bCFNs. In addition, we expressed a GFP-Gephyrin fusion protein in bCFNs using the transgenic line [*Et(REX2-SCP1:GAL4FF)y321;Tg(UAS:Geph-GFP)*]. In this labeling, we found that Syt2a puncta colocalized with postsynaptic GFP-Geph puncta on bCFN neurites (Fig.4F-F”’), suggesting that Syt2a is localized in presynaptic terminals formed onto projections from bCFNs at (but not exclusively) inhibitory synapses. From these experiments, we conclude that Syt2a is both expressed at presynaptic terminals from bCFNs, and that Syt2a is localized at presynaptic terminals that synapse onto bCFN somata and neurites.

### Syt2a and tyrosine hydroxylase colocalize at synapses

Dopaminergic neurons in the tegmentum that project anteriorly to ventral forebrain regions, have been identified as important regulators of social behaviors in vertebrates (Arakawa and Ikeda, 1991). Tyrosine hydroxylase (TH) is the rate limiting enzyme for catecholamine synthesis and is necessary for dopamine production. There are two *th* genes in zebrafish, *th1* and *th2*, with *th1* being expressed in the olfactory bulb, telencephalon, preoptic area, posterior tuberculum, hypothalamus and hindbrain, while *th2* is only expressed in the diencephalon and midbrain (Filippi et al., 2010; Yamamoto et al., 2010). To label all TH-positive neurons, we used an antibody that recognizes zebrafish TH1 and TH2 proteins (Yamamoto et al., 2010). We found TH labeling in cell bodies and projections in many brain regions (Fig.5A-C), similar to TH distribution in the adult zebrafish brain (Rink and Wullimann, 2002; Yamamoto et al., 2010). In general, TH labeling mostly reflected the *th1* mRNA expression pattern (Filippi et al., 2010). TH antibody labels neurons in the olfactory bulb, dorsal and ventral telencephalon, preoptic area, hypothalamus, thalamus and hindbrain (Fig.5B,C). We did not detect TH-positive cell bodies in the optic tectum, but found TH-positive neurites in two bands located within the retino-recipient strata and deeper strata of the optic tectum neuropil (Fig.5C, arrow) (Arenzana et al., 2006). The ventral telencephalon has only a few TH expressing neurons, but we expected to see TH-positive synaptic puncta in the medAC and in the region adjacent to the medAC (Fig.5D-D”), as suggested by previous studies that traced the projections of dopaminergic neurons (Du et al., 2016). Indeed, we found TH puncta in and adjacent to the medAC, and these puncta colocalized with Syt2a, suggesting that TH is located in presynaptic terminals (Fig.5D-D”). Further, we found TH positive cell bodies distributed throughout the olfactory bulb and punctate TH labeling on olfactory bulb projections (Fig.5E-E”’). These results confirm that TH puncta are localized on projections and even presynaptic terminals, and further suggest a localization of TH to Syt2a-positive terminals in the ventral telencephalon.

**Fig.5.**
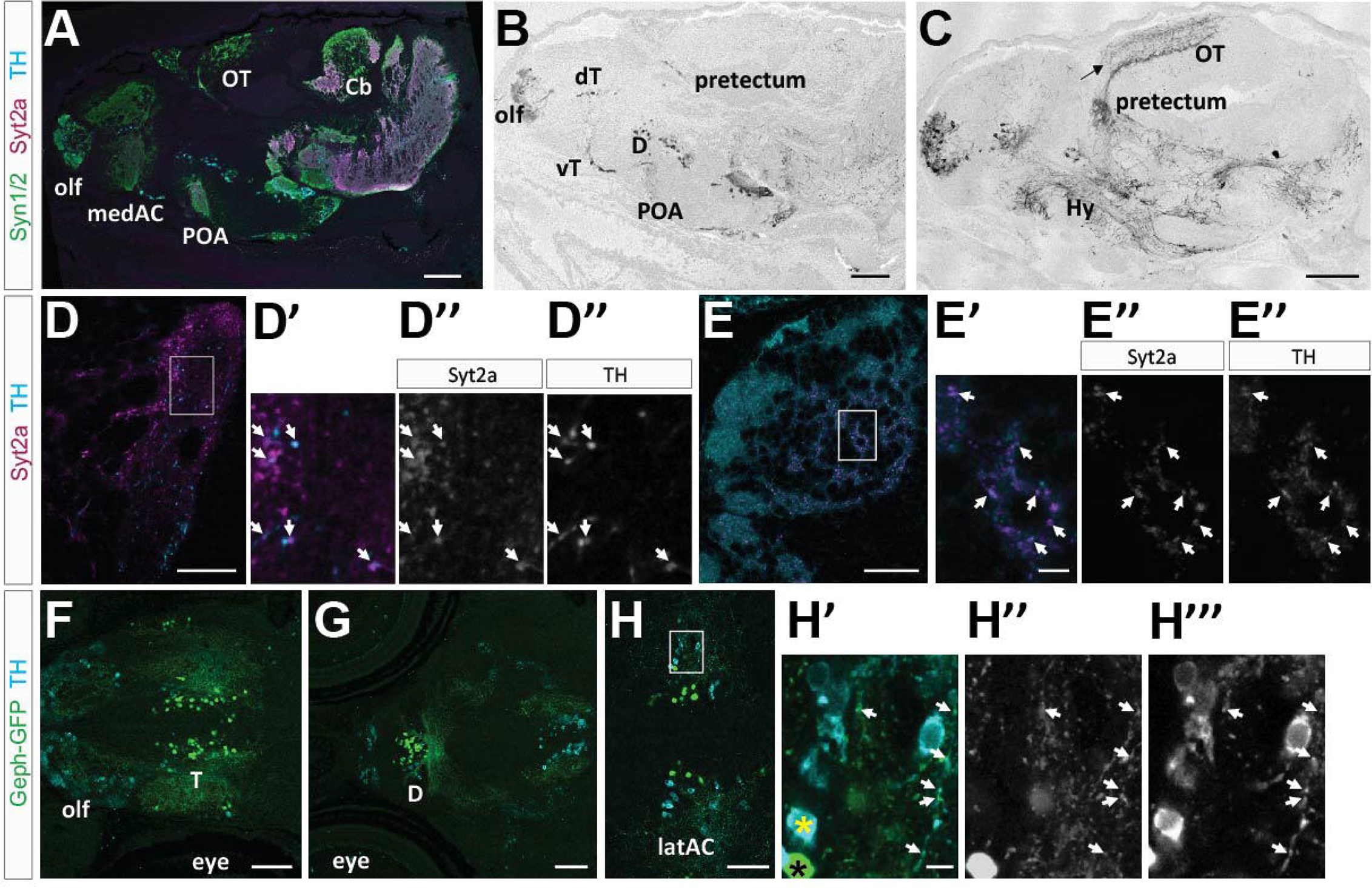
Tyrosine hydroxylase labeling and co-localization with Syt2a. **5A-C**. Sagittal view of a brain at 14 dpf, showing TH labeling in distinct neuronal clusters throughout the brain. **5A.** Labeling showing TH (cyan), Syt2a (magenta) and Syn1/2 (green). **5B,C**. TH labels neurons in the olfactory bulb (olf), dorsal (dT) and ventral telencephalon (vT), diencephalon (D), preoptic area (POA) and pretectum. TH projections are located throughout the brain. An arrow marks labeling in the optic tectum (OT). **5D-D”**. Syt2a and TH co-localize at synapses in the AC. Arrows mark Syt2a (D’) and TH (D”) co-labeled puncta. **5E-E”**. In the olfactory bulb we find many Syt2 (E’) and TH (E”) co-labeled puncta (arrows). **5F-H”’**. Dorsal view of the forebrain showing TH labeling (cyan) and Geph-GFP positive (green) specifically in bCFNs. **5F**. Labeling in the olf and telencephalon (T). **5G**. More posterior view showing TH labeling and Geph-GFP positive bCFNs in the diencephalon (D). **5H-H”’**. Higher magnification of the telencephalic region. A yellow and black asterisk marks TH only and Geph-GFP only neurons, respectively. Arrows point to TH and Geph-GFP co-localization on projections from bCFNs. Area shown in 5H’-H”’ is marked with a box in 5H. Scale bars are 200 μm in A-C, 30 μm in D,E, 70 μm in F,G, 50 μm in H and 5 μm in H’.

Because dopamine is important for social behavior (Arakawa and Ikeda, 1991; Stednitz et al., 2018), we explored whether TH puncta are located on forebrain bCFN cell bodies or their neurites. We did not detect TH in bCFNs cell bodies (Fig.5H-H”’; TH, black asterisk; bCFN, yellow asterisk), but found that TH-positive neuronal cell bodies are often located in the vicinity of GFP-Geph expressing bCFNs (5F-H). Further, TH puncta frequently colocalized with GFP-Geph puncta (arrows in Fig. 5H-H”’), suggesting that inhibitory presynaptic terminals, located on the neurites of bCFNs, are also TH-positive. These studies suggest that Syt2a protein localizes to synapses in a region of the brain and at a time in development that is associated with the initiation of robust social behavior. Furthermore, Syt2a-positive synapses impinging on the neurites of bCFNs in the ventral telencephalon show TH labeling, suggesting that they might release dopamine, a neurotransmitter associated with social behavior.

## DISCUSSION

Among the many Synaptotagmins described, Syt1 and Syt2 are the major proteins mediating fast and synchronous vesicular release at central synapses (Fernández-Chacón et al., 2001; Pang et al., 2006a; Pang et al., 2006b). Despite the similarity between Syt1 and Syt2, and their sometimes redundant function, Syt2 release kinetics are substantially faster than release triggered by Syt1 (Xu et al., 2007), and thus differential expression of Syt1 and Syt2 has the potential to significantly alter the function of synapses during development. Here we have examined the expression pattern of one of the Syt2 homologs in the zebrafish brain during development, with a focus on brain regions implicated in social behavior.

We found that zebrafish Syt2a protein is highly expressed in the midbrain and hindbrain at 6 dpf but in contrast, Syt2 expression in the forebrain was very low. Interestingly, we found that Syt2a distribution in the forebrain changes over an identified developmental period. Syt2 expression in the forebrain at 6 dpf steadily increases until it shows robust expression at 14 dpf, a time at which zebrafish larvae start to show complex behaviors, including social behavior (Dreosti et al., 2015; Larsch and Baier, 2018). Although the late onset of Syt2a expression in the forebrain could be explained by ascending projections from more posterior regions that eventually grow into the forebrain, we favor a second explanation. Our data suggest that Syt2a protein is generated at low levels in the forebrain until about 14 dpf, when Syt2a protein synthesis is dramatically upregulated. This is supported by the presence of *syt2a* transcript in the forebrain at 6 dpf (suppl Fig). Our data suggest that already existing synapses in the forebrain change their protein composition and potentially also their electrophysiological properties as part of a synaptic refinement step around 14 dpf, the time at which social behavior becomes robust (Dreosti et al., 2015; Larsch and Baier, 2018).

In mice, Syt2 is widely expressed in the spinal cord, brainstem, and cerebellum at P15. In addition, similar to zebrafish, Syt2 is present in selected forebrain neurons, including most striatal neurons and some hypothalamic, cortical, and hippocampal neurons (Fox and Sanes, 2007; Pang et al., 2006a). However, the expression pattern of mouse Syt2 has not been extensivily described at different developmental timepoints, and it is therefore unclear whether regions of the mouse brain might dramatically increase Syt2 expression over the course of development.

It is intriguing that the onset of Syt2a expression in the forebrain and the onset of robust social behavior coincide at around 14 dpf, implicating that Syt2a is involved in the functional refinement of circuits by modulating calcium sensitivity and vesicle release properties of previously established synapses. We observed a significant increase in Syt2a protein expression in the ventral telencephalon, specifically in the AC and in regions adjacent to the AC associating with bCFNs. In previous work, we showed that this specific population of cholinergic neurons in the ventral forebrain is required for social behavior in adult zebrafish (Stednitz et al., 2018). Here, we show, that Syt2a is located at synapses onto somata of bCFNs and also in their neurites, suggesting that Syt2a is important in this circuit and might define the developmental time line of social behavior acquisition. These findings are consistent with a suggested social behavior network (SBN, Geng and Peterson, 2019), consisting of several identified regions within the mammalian forebrain. Despite the lack of some mammalian-specific regions, the zebrafish forebrain shares most regions implicated in the mammalian SBN suggesting that this network is highly conserved (Geng and Peterson, 2019). Different nomenclature in mammals and zebrafish complicates the direct anatomical comparison. However, the mammalian nucleus accumbens, lateral septum, striatum and medial preoptic regions are homologous regions to the zebrafish ventral telencephalon and the preoptic region, both regions in which we find Syt2a protein at the time when social behavior is strengthened. The neurons upstream and downstream of bCFNs have not been identified; a crucial step to fully elucidating this social behavioral circuit.

Studies in mouse, rat and zebrafish have shown that dopamine is essential for social behavior (Arakawa and Ikeda, 1991; Homberg et al., 2016; Oliveri and Levin, 2019; Parker et al., 2013; Stednitz et al., 2018). We asked whether dopaminergic input might impinge upon the bCFNs in the ventral telencephalon, by examining TH expression pattern at 14 dpf. TH is a major biosynthetic enzyme for dopamine and other catecholamines. Our experiments revealed several important observations: (1) bCFNs in the ventral telencephalon do not express TH, (2) TH puncta colocalize with Syt2a and GFP-Geph on neurites from bCFNs, and (3) there are several clusters of TH-positive neurons that might be sources of TH-positive presynaptic terminals onto bCFNs in the ventral telencephalon.

The finding of an association between TH and Syt2a is consistent with findings in mammals, which showed that Syt2 colocalizes with TH mRNA in axons of dissociated superior cervical ganglion neurons. This study demonstrated that TH is locally synthesized and that TH protein accumulates with Synaptotagmin in presynaptic terminals (Gervasi et al., 2016), analogous to our results. Furthermore, we identified numerous clusters of neurons expressing TH, some of which had obvious projections to the ventral telencephalon. These include the olfactory bulb, the ventral telencephalon and the diencephalon. Similar populations have been observed in mice, suggesting that homologous populations of dopaminergic neurons might be participating in the regulation of social behavior in mammals.

TH-expressing neurons are mostly located in the hypothalamus, striatum and cortex in mammals. The ventral tegmental area in the midbrain is one of the major sources of mesostriatal dopaminergic projections in mammals, but not in zebrafish (Rink and Wullimann, 2002). Neurons in the ventral tegmentum project into multiple limbic forebrain areas, including the nucleus accumbens, located in the basal forebrain rostral to the preoptic region of the hypothalamus, a component of the social behavior network (Alger et al., 2011). This distribution is largely conserved among vertebrates (Smeets and González, 2000). However, zebrafish lack dopaminergic neurons in the mesencephalon (Filippi et al., 2010; Rink and Wullimann, 2001), and the zebrafish diencephalic TH-positive cluster has been assumed to be a functionally homologous structure (Rink and Wullimann, 2002; Tay et al., 2011; Yamamoto and Vernier, 2011). Although it is possible that the projections onto the bCFNs come from any of the TH-positive clusters, we hypothesize that the ventral diencephalic cluster is the source of this input.

In the future, it will be interesting to test whether brain region-specific deletion of Syt2 will significantly alter behaviors in the zebrafish, whether they manifest at a time point at which we first detect Syt2 protein at specific synapses on bCFNs in the ventral telencephalon, and whether dopaminergic input into the ventral diencephalon is altered by this genetic manipulation.

## MATERIALS AND METHODS

### Zebrafish husbandry and lines

Zebrafish embryos for the 4tU-labeling experiments were obtained from natural spawning of wild-type (AB or AB/TU) or the transgenic lines *Et(REX2-SCP1:GAL4FF)y321* (Marquart et al., 2015)*, Tg(UAS:GFP) (Kawakami et al., 2016), Tg(UAS:Syn1-GFP)* (Easley-Neal et al., 2013), *Tg(UAS:GFP-Geph)* in the University of Oregon Zebrafish Facility, and were staged at 28.5°C accordingly (Kimmel et al., 1994). All procedures were carried out under an approved protocol with the University of Oregon Animal Care and Use Committee.

To generate the transgenic line *Tg(UAS:GFP-Geph)*, purified DNA was injected into fertilized wild-type (AB) zebrafish eggs at the one-cell stage together with RNA encoding Tol2 transposase, following standard protocols.

### Syt2 homologs in the zebrafish

Zebrafish Synaptotagmin 2 (Syt2) homologs are highly similar in their amino acid composition. The *syt2a* (ZDB-GENE-060503-315) and *syt2b* (ZDB-GENE-090717-1) duplicates are located on Chromosomes 23 and 6 (Sanger zebrafish reference genome assembly GRCz11), respectively (Table 1). A knock-down of Syt2 (chr.6) leads to loss of znp-1 antibody labeling, showing that znp-1 selectively recognizes Syt2 [chr.6, (Wen et al., 2010)]. However, because of the location of the gene *syt2a* on chr.23 in an environment that exclusively harbors b duplicates, *syt2a* was renamed to *syt2b* (Liu et al., 2015).

**Table 1.** Gene reference sources.

| gene name | e!Ensembl location | NCBI Reference Sequence ID | Sanger Gene ID | ZFIN Gene ID | antibody |
| --- | --- | --- | --- | --- | --- |
| *syt2a* | chromosome 6:<br>55174657-55197620 | XM_6830000.7 | not annotated | ZDB-GENE-090717-1 | znp-1 |
| *syt2b* | chromosome<br>23: 6,232,895-6,313,808 | XM_005174112.4 | ENSDARG00000025206 | ZDB-GENE-060503-315 | None |

### Cloning

We used the primers syt2a-F2:5’-ACAACTCCACCGAGTCTGAG-3’ and syt2a-R1: 5’- ATACACAGACATGACCAGCG -3’ to clone a 592 bp fragment, amplified from zebrafish cDNA, into pCR-Blunt II-TOPO-vector (ThermoFisher).

The construct *UAS:GFP-Geph* was generated using the Gateway system. The cDNA encoding zebrafish Gephyrinb (a gift from Robert Harvey) was cloned into a p3E entry vector and combined with *p5E-4xnr:UAS* and *pME-GFP* into a destination vector.

### Immunohistochemistry

RNA in situ hybridization and immunohistochemistry were carried out on brain cryostat sections (16 um) according to standard protocols (Westerfield, 2000). To optimize labeling of the anti-GluR 2/3 antibody, we used a “fresh-frozen” protocol entailing cryosectioning on non-fixed, frozen embryos. Cryosectioned tissues underwent a post-sectioning fixation in 4% PFA-PBS solution for eight minutes.

For immunohistochemistry the following primary antibodies were used: Syn1/2 (rb, Synaptic Systems) at 1:250, SV2 (ms, Developmental Studies Hybridoma Bank) at 1:1000, znp-1 (Syt2a, IgG2a, Zebrafish International Resource Center) at 1:500, Gephyrin (rb, Abcam) at 1:1000, GluR2/3 (IgG2a, Millipore Sigma) at 1:250, TH (rb, Millipore Sigma) at 1: 500, GFP (ch, Aves Labs) at 1:1000. Primary antibodies were revealed using the following secondary antibodies at a concentration of 1:750 [Alexa Fluor goat anti-mouse IgG1, IgG_2a_, and (H+L), Molecular Probes, coupled to 488, 546 or 633], Alexa Fluor 633 goat anti-rabbit, Molecular Probes, Alexa Fluor 488 goat anti-chicken, Molecular Probes and goat anti-rabbit Cy5 [IgG (H+L), Jackson ImmunoResearch Laboratories].

### Confocal Microscopy

Images were taken on an inverted Nikon TU-2000 microscope with an EZ-C1 confocal system (Nikon) with either a 20x, 60x water immersion, or 100x oil immersion objective, a Zeiss LSM5 Pascal confocal microscope with ZEN software with 20x or 63x oil immersion objectives or a Leica SP8 confocal microscope with LasX software with 40x water or 63x oil immersion objectives.

### Quantification of synaptic puncta

Cropped sections of the 100x or 63x images were saved as separate channels in grayscale bitmap format. Using the Image Pro Plus® (Media Cybernetics) program, puncta labeled by Synapsin 1/2, Gephyrin, and GluR 2/3 were counted manually and co-localization was quantified as performed previously (Hoy et al., 2013). Intensity was measured in cropped sections of 20x images using ImageJ and plotted as neuropil intensity (arbitrary unit, a.u.).

## REFERENCES

Alger, S.J., Juang, C., Riters, L.V., 2011. Social affiliation relates to tyrosine hydroxylase immunolabeling in male and female zebra finches (Taeniopygia guttata). Journal of chemical neuroanatomy 42, 45–55.

Arakawa, O., Ikeda, T., 1991. Apomorphine effect on single and paired rat open-field behavior. Physiology & Behavior 50, 189–194.

Arenzana, F.J., Arévalo, R., Sánchez-González, R., Clemente, D., Aijón, J., Porteros, A., 2006. Tyrosine hydroxylase immunoreactivity in the developing visual pathway of the zebrafish. Anatomy and Embryology 211, 323.

Bouhours, B., Gjoni, E., Kochubey, O., Schneggenburger, R., 2017. Synaptotagmin2 (Syt2) Drives Fast Release Redundantly with Syt1 at the Output Synapses of Parvalbumin-Expressing Inhibitory Neurons. The Journal of Neuroscience 37, 4604.

Chen, C., Arai, I., Satterfield, R., Young, S.M., Jr., Jonas, P., 2017. Synaptotagmin 2 Is the Fast Ca(2+) Sensor at a Central Inhibitory Synapse. Cell reports 18, 723–736.

Dreosti, E., Lopes, G., Kampff, A.R., Wilson, S.W., 2015. Development of social behavior in young zebrafish. Frontiers in neural circuits 9, 39–39.

Du, Y., Guo, Q., Shan, M., Wu, Y., Huang, S., Zhao, H., Hong, H., Yang, M., Yang, X., Ren, L., Peng, J., Sun, J., Zhou, H., Li, S., Su, B., 2016. Spatial and Temporal Distribution of Dopaminergic Neurons during Development in Zebrafish. Frontiers in neuroanatomy 10, 115–115.

Easley-Neal, C., Fierro, J., Jr., Buchanan, J., Washbourne, P., 2013. Late recruitment of synapsin to nascent synapses is regulated by Cdk5. Cell Rep 3, 1199–1212.

Engeszer, R.E., Barbiano, L.A.D.A., Ryan, M.J., Parichy, D.M., 2007. Timing and plasticity of shoaling behaviour in the zebrafish, Danio rerio. Animal behaviour 74, 1269–1275.

Fernández-Chacón, R., Königstorfer, A., Gerber, S.H., García, J., Matos, M.F., Stevens, C.F., Brose, N., Rizo, J., Rosenmund, C., Südhof, T.C., 2001. Synaptotagmin I functions as a calcium regulator of release probability. Nature 410, 41–49.

Filippi, A., Mahler, J., Schweitzer, J., Driever, W., 2010. Expression of the paralogous tyrosine hydroxylase encoding genes th1 and th2 reveals the full complement of dopaminergic and noradrenergic neurons in zebrafish larval and juvenile brain. The Journal of comparative neurology 518, 423–438.

Fox, M.A., Sanes, J.R., 2007. Synaptotagmin I and II are present in distinct subsets of central synapses. Journal of Comparative Neurology 503, 280–296.

Garner, C.C., Waites, C.L., Ziv, N.E., 2006. Synapse development: still looking for the forest, still lost in the trees. Cell and Tissue Research 326, 249.

Geng, Y., Peterson, R.T., 2019. The zebrafish subcortical social brain as a model for studying social behavior disorders. Dis Model Mech 12.

Gervasi, N.M., Scott, S.S., Aschrafi, A., Gale, J., Vohra, S.N., MacGibeny, M.A., Kar, A.N., Gioio, A.E., Kaplan, B.B., 2016. The local expression and trafficking of tyrosine hydroxylase mRNA in the axons of sympathetic neurons. RNA 22, 883–895.

Homberg, J.R., Olivier, J.D., VandenBroeke, M., Youn, J., Ellenbroek, A.K., Karel, P., Shan, L., van Boxtel, R., Ooms, S., Balemans, M., Langedijk, J., Muller, M., Vriend, G., Cools, A.R., Cuppen, E., Ellenbroek, B.A., 2016. The role of the dopamine D1 receptor in social cognition: studies using a novel genetic rat model. Dis Model Mech 9, 1147–1158.

Hoy, J.L., Haeger, P.A., Constable, J.R., Arias, R.J., McCallum, R., Kyweriga, M., Davis, L., Schnell, E., Wehr, M., Castillo, P.E., Washbourne, P., 2013. Neuroligin1 drives synaptic and behavioral maturation through intracellular interactions. J Neurosci 33, 9364–9384.

Kawakami, K., Asakawa, K., Hibi, M., Itoh, M., Muto, A., Wada, H., 2016. Gal4 Driver Transgenic Zebrafish: Powerful Tools to Study Developmental Biology, Organogenesis, and Neuroscience. Adv Genet 95, 65–87.

Kimmel, C.B., Warga, R.M., Kane, D.A., 1994. Cell cycles and clonal strings during formation of the zebrafish central nervous system. Development 120, 265.

Kochubey, O., Babai, N., Schneggenburger, R., 2016. A Synaptotagmin Isoform Switch during the Development of an Identified CNS Synapse. Neuron 91, 1183.

Larsch, J., Baier, H., 2018. Biological Motion as an Innate Perceptual Mechanism Driving Social Affiliation. Current Biology 28, 3523–3532.e3524.

Liu, C., Hu, J., Qu, C., Wang, L., Huang, G., Niu, P., Zhong, Z., Hong, F., Wang, G., Postlethwait, J.H., Wang, H., 2015. Molecular evolution and functional divergence of zebrafish (Danio rerio) cryptochrome genes. Scientific Reports 5, 8113.

Marquart, G.D., Tabor, K.M., Brown, M., Strykowski, J.L., Varshney, G.K., LaFave, M.C., Mueller, T., Burgess, S.M., Higashijima, S.-I., Burgess, H.A., 2015. A 3D Searchable Database of Transgenic Zebrafish Gal4 and Cre Lines for Functional Neuroanatomy Studies. Frontiers in neural circuits 9, 78–78.

O’Connell, L.A., Hofmann, H.A., 2012. Evolution of a vertebrate social decision-making network. Science 336, 1154–1157.

Oliveri, A.N., Levin, E.D., 2019. Dopamine D1 and D2 receptor antagonism during development alters later behavior in zebrafish. Behavioural Brain Research 356, 250–256.

Pang, Z.P., Melicoff, E., Padgett, D., Liu, Y., Teich, A.F., Dickey, B.F., Lin, W., Adachi, R., Südhof, T.C., 2006a. Synaptotagmin-2 Is Essential for Survival and Contributes to Ca^2+^ Triggering of Neurotransmitter Release in Central and Neuromuscular Synapses. The Journal of Neuroscience 26, 13493.

Pang, Z.P., Sun, J., Rizo, J., Maximov, A., Südhof, T.C., 2006b. Genetic analysis of synaptotagmin 2 in spontaneous and Ca2+-triggered neurotransmitter release. The EMBO journal 25, 2039–2050.

Parker, K.L., Alberico, S.L., Miller, A.D., Narayanan, N.S., 2013. Prefrontal D1 dopamine signaling is necessary for temporal expectation during reaction time performance. Neuroscience 255, 246–254.

Rink, E., Wullimann, M.F., 2001. The teleostean (zebrafish) dopaminergic system ascending to the subpallium (striatum) is located in the basal diencephalon (posterior tuberculum). Brain Research 889, 316–330.

Rink, E., Wullimann, M.F., 2002. Connections of the ventral telencephalon and tyrosine hydroxylase distribution in the zebrafish brain (Danio rerio) lead to identification of an ascending dopaminergic system in a teleost. Brain Research Bulletin 57, 385–387.

Rizo, J., Xu, J., 2015. The Synaptic Vesicle Release Machinery. Annual Review of Biophysics 44, 339–367.

Roberts, A.C., Bill, B.R., Glanzman, D.L., 2013. Learning and memory in zebrafish larvae. Frontiers in neural circuits 7, 126–126.

Smeets, W.J.A.J., González, A., 2000. Catecholamine systems in the brain of vertebrates: new perspectives through a comparative approach. Brain Research Reviews 33, 308–379.

Stednitz, S.J., McDermott, E.M., Ncube, D., Tallafuss, A., Eisen, J.S., Washbourne, P., 2018. Forebrain Control of Behaviorally Driven Social Orienting in Zebrafish. Curr Biol. 28, 2445–2451.e3.

Südhof, T.C., 2002. Synaptotagmins: Why So Many? Journal of Biological Chemistry 277, 7629–7632.

Tay, T.L., Ronneberger, O., Ryu, S., Nitschke, R., Driever, W., 2011. Comprehensive catecholaminergic projectome analysis reveals single-neuron integration of zebrafish ascending and descending dopaminergic systems. Nature communications 2, 171–171.

Wen, H., Linhoff, M.W., McGinley, M.J., Li, G.-L., Corson, G.M., Mandel, G., Brehm, P., 2010. Distinct roles for two synaptotagmin isoforms in synchronous and asynchronous transmitter release at zebrafish neuromuscular junction. Proceedings of the National Academy of Sciences 107, 13906.

Westerfield, M., 2000. The Zebrafish Book, 4th ed. Institute for Neuroscience, University of Oregon, Eugene OR.

Xu, J., Mashimo, T., Südhof, T.C., 2007. Synaptotagmin-1, −2, and −9: Ca2+ Sensors for Fast Release that Specify Distinct Presynaptic Properties in Subsets of Neurons. Neuron 54, 567–581.

Yamamoto, K., Ruuskanen, J.O., Wullimann, M.F., Vernier, P., 2010. Two tyrosine hydroxylase genes in vertebrates: New dopaminergic territories revealed in the zebrafish brain. Molecular and Cellular Neuroscience 43, 394–402.

Yamamoto, K., Vernier, P., 2011. The Evolution of Dopamine Systems in Chordates. Frontiers in Neuroanatomy 5.

Zhao, Y., Marin, O., Hermesz, E., Powell, A., Flames, N., Palkovits, M., Rubenstein, J.L., Westphal, H., 2003. The LIM-homeobox gene Lhx8 is required for the development of many cholinergic neurons in the mouse forebrain. Proc Natl Acad Sci U S A 100, 9005–9010.

