## Supplementary figures and images for "Late onset of Syt2a expression at synapses relevant to social behavior"

### Supplemental Fig. 1

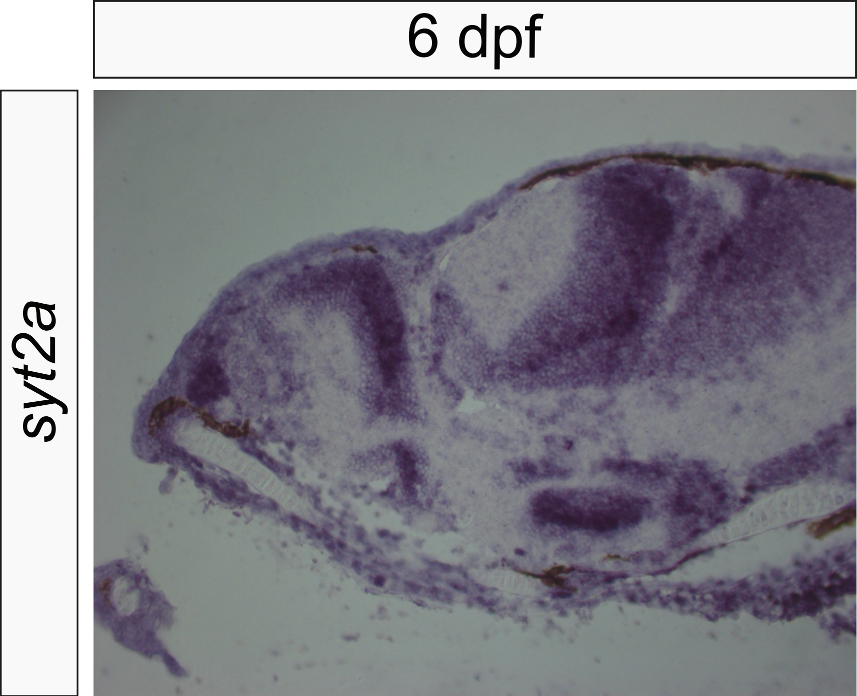
